# Single-cell transcriptomic characterization of 20 organs and tissues from individual mice creates a *Tabula Muris*

**DOI:** 10.1101/237446

**Authors:** Nicholas Schaum, Jim Karkanias, Norma F Neff, Andrew P. May, Stephen R. Quake, Tony Wyss-Coray, Spyros Darmanis, Joshua Batson, Olga Botvinnik, Michelle B. Chen, Steven Chen, Foad Green, Robert Jones, Ashley Maynard, Lolita Penland, Rene V. Sit, Geoffrey M. Stanley, James T. Webber, Fabio Zanini, Ankit S. Baghel, Isaac Bakerman, Ishita Bansal, Daniela Berdnik, Biter Bilen, Douglas Brownfield, Corey Cain, Michelle B. Chen, Steven Chen, Min Cho, Giana Cirolia, Stephanie D. Conley, Spyros Darmanis, Aaron Demers, Kubilay Demir, Antoine de Morree, Tessa Divita, Haley du Bois, Laughing Bear Torrez Dulgeroff, Hamid Ebadi, F. Hernan Espinoza, Matt Fish, Qiang Gan, Benson M. George, Astrid Gillich, Foad Green, Geraldine Genetiano, Xueying Gu, Gunsagar S. Gulati, Yan Hang, Shayan Hosseinzadeh, Albin Huang, Tal Iram, Taichi Isobe, Feather Ives, Robert Jones, Kevin S. Kao, Guruswamy Karnam, Aaron M. Kershner, Bernhard Kiss, William Kong, Maya E. Kumar, Jonathan Lam, Davis P. Lee, Song E. Lee, Guang Li, Qingyun Li, Ling Liu, Annie Lo, Wan-Jin Lu, Anoop Manjunath, Andrew P. May, Kaia L. May, Oliver L. May, Ashley Maynard, Marina McKay, Ross J. Metzger, Marco Mignardi, Dullei Min, Ahmad N. Nabhan, Norma F Neff, Katharine M. Ng, Joseph Noh, Rasika Patkar, Weng Chuan Peng, Lolita Penland, Robert Puccinelli, Eric J. Rulifson, Nicholas Schaum, Shaheen S. Sikandar, Rahul Sinha, Rene V Sit, Krzysztof Szade, Weilun Tan, Cristina Tato, Krissie Tellez, Kyle J. Travaglini, Carolina Tropini, Lucas Waldburger, Linda J. van Weele, Michael N. Wosczyna, Jinyi Xiang, Soso Xue, Justin Youngyunpipatkul, Fabio Zanini, Macy E. Zardeneta, Fan Zhang, Lu Zhou, Ishita Bansal, Steven Chen, Min Cho, Giana Cirolia, Spyros Darmanis, Aaron Demers, Tessa Divita, Hamid Ebadi, Geraldine Genetiano, Foad Green, Shayan Hosseinzadeh, Feather Ives, Annie Lo, Andrew P. May, Ashley Maynard, Marina McKay, Norma F. Neff, Lolita Penland, Rene V. Sit, Weilun Tan, Lucas Waldburger, Justin Youngyunpipatkul, Joshua Batson, Olga Botvinnik, Paola Castro, Derek Croote, Spyros Darmanis, Joseph L. DeRisi, Jim Karkanias, Angela Pisco, Geoffrey M. Stanley, James T. Webber, Fabio Zanini, Ankit S. Baghel, Isaac Bakerman, Joshua Batson, Biter Bilen, Olga Botvinnik, Douglas Brownfield, Michelle B. Chen, Spyros Darmanis, Kubilay Demir, Antoine de Morree, Hamid Ebadi, F. Hernan Espinoza, Matt Fish, Qiang Gan, Benson M. George, Astrid Gillich, Xueying Gu, Gunsagar S. Gulati, Yan Hang, Albin Huang, Tal Iram, Taichi Isobe, Guruswamy Karnam, Aaron M. Kershner, Bernhard M. Kiss, William Kong, Christin S. Kuo, Jonathan Lam, Benoit Lehallier, Guang Li, Qingyun Li, Ling Liu, Wan-Jin Lu, Dullei Min, Ahmad N. Nabhan, Katharine M. Ng, Patricia K. Nguyen, Rasika Patkar, Weng Chuan Peng, Lolita Penland, Eric J. Rulifson, Nicholas Schaum, Shaheen S. Sikandar, Rahul Sinha, Krzysztof Szade, Serena Y. Tan, Krissie Tellez, Kyle J. Travaglini, Carolina Tropini, Linda J. van Weele, Bruce M. Wang, Michael N. Wosczyna, Jinyi Xiang, Hanadie Yousef, Lu Zhou, Joshua Batson, Olga Botvinnik, Steven Chen, Spyros Darmanis, Foad Green, Andrew P. May, Ashley Maynard, Angela Pisco, Stephen R. Quake, Nicholas Schaum, Geoffrey M. Stanley, James T. Webber, Tony Wyss-Coray, Fabio Zanini, Philip A. Beachy, Charles K. F. Chan, Antoine de Morree, Benson M. George, Gunsagar S. Gulati, Yan Hang, Kerwyn Casey Huang, Tal Iram, Taichi Isobe, Aaron M. Kershner, Bernhard M. Kiss, William Kong, Guang Li, Qingyun Li, Ling Liu, Wan-Jin Lu, Ahmad N. Nabhan, Katharine M. Ng, Patricia K. Nguyen, Nicholas Schaum, Shaheen S. Sikandar, Rahul Sinha, Krzysztof Szade, Kyle J. Travaglini, Carolina Tropini, Bruce M. Wang, Kenneth Weinberg, Michael N. Wosczyna, Sean Wu, Hanadie Yousef, Ben A. Barres, Philip A. Beachy, Charles K. F. Chan, Michael F. Clarke, Spyros Darmanis, Kerwyn Casey Huang, Jim Karkanias, Seung K. Kim, Mark A. Krasnow, Christin S. Kuo, Andrew P. May, Norma Neff, Roel Nusse, Patricia K. Nguyen, Thomas A. Rando, Justin Sonnenburg, Bruce M. Wang, Kenneth Weinberg, Irving L. Weissman, Sean M. Wu, Stephen R. Quake, Tony Wyss-Coray

## Abstract

We have created a compendium of single cell transcriptome data from the model organism *Mus musculus* comprising more than 100,000 cells from 20 organs and tissues. These data represent a new resource for cell biology, revealing gene expression in poorly characterized cell populations and allowing for direct and controlled comparison of gene expression in cell types shared between tissues, such as T-lymphocytes and endothelial cells from distinct anatomical locations. Two distinct technical approaches were used for most tissues: one approach, microfluidic droplet-based 3’-end counting, enabled the survey of thousands of cells at relatively low coverage, while the other, FACS-based full length transcript analysis, enabled characterization of cell types with high sensitivity and coverage. The cumulative data provide the foundation for an atlas of transcriptomic cell biology.

---

The cell is a fundamental unit of structure and function in biology, and multicellular organisms have evolved a wide variety of different cell types with specialized roles. Although cell types have historically been characterized on the basis of morphology and phenotype, the development of molecular methods has enabled ever more precise defining of their properties, typically by measuring protein or mRNA expression patterns^1^. Technological advances have enabled increasingly greater degrees of multiplexing of these measurements^2-7^, and it is now possible to use highly parallel sequencing to enumerate nearly every mRNA molecule in a given single cell^7,8^. This approach has provided many novel insights into cell biology and the composition of organs from a variety of organisms^9-18^. However, while these reports provide valuable characterization of individual organs, it is challenging to compare data taken with varying experimental techniques in independent labs from different animals. It therefore remains an open question whether data from individual organs can be synthesized and used as a more general resource for biology.

Here we report a compendium of cell types from the mouse *Mus musculus.* We analyzed multiple organs and tissues from the same animal, thereby generating a data set controlled for age, environment and epigenetic effects. This enables the direct comparison of cell type composition between organs as well as comparison of shared cell types across the entire organism. The compendium is comprised of single cell transcriptome sequence data from 100,605 cells isolated from 20 organs and tissues (Fig. 1). Those were collected from 3 female and 4 male, C57BL/6 NIA, 3 month old mice (10-15 weeks), whose developmental age is roughly analogous to humans at 20 years of age. All data, protocols, and analysis scripts from the *Tabula Muris* are shared as a public resource (http://tabula-muris.ds.czbiohub.org/), gene counts and metadata from all single cells are accessible on Figshare (https://figshare.com/account/home#/projects/27733), raw data are available on GEO (GSE109774), and all code used for analysis is available on GitHub (https://github.com/czbiohub/tabula-muris). While these data are by no means a complete representation of all mouse organs and cell types, they provide a first draft attempt to create an organism-wide representation of cellular diversity and a comparative framework for future studies using the large variety of murine disease models.

**Figure 1.**
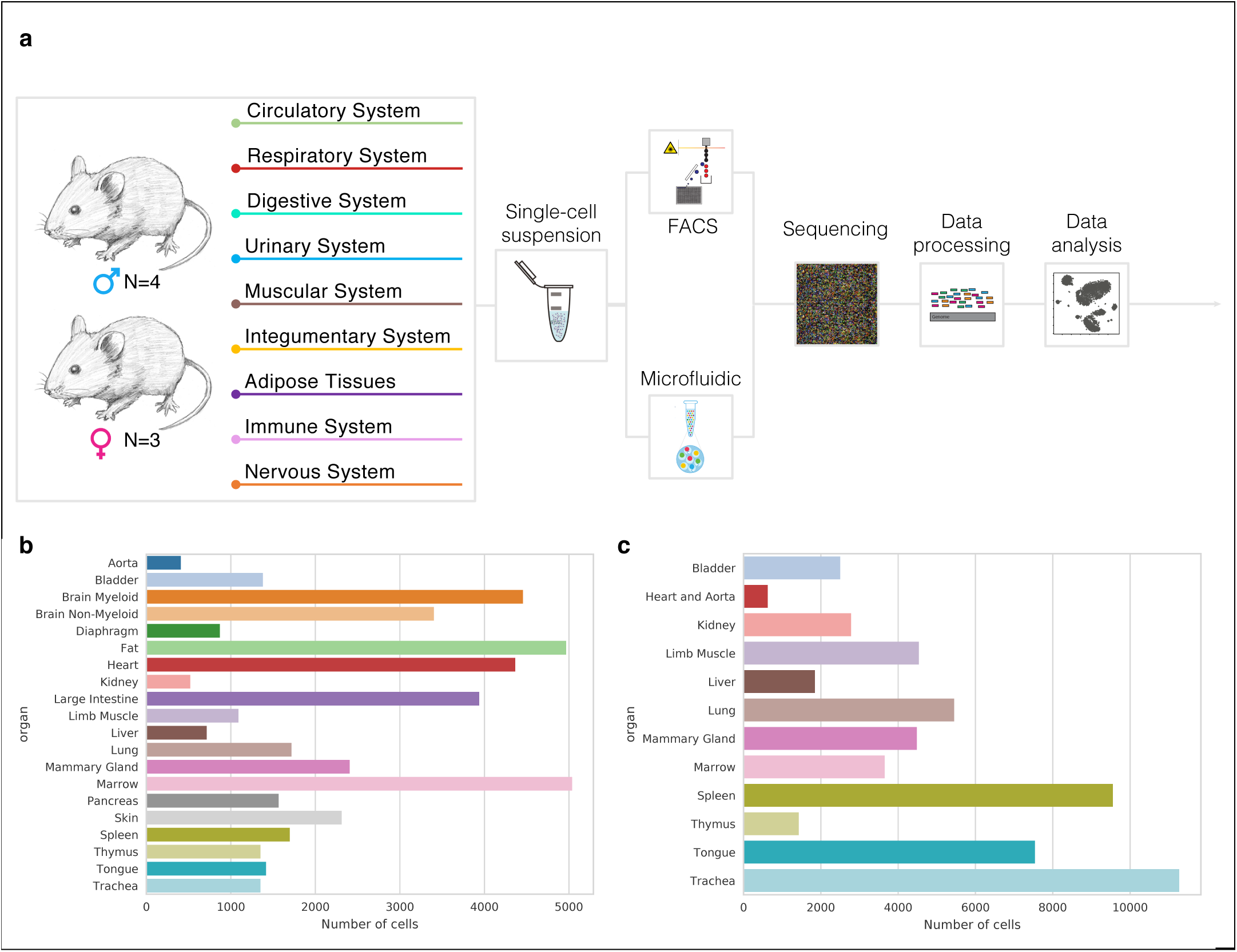
Overview of *Tabula Muris*. a) 20 organs and tissues from 4 male and 3 female mice were analyzed. After dissociation, cells were either sorted by FACS or captured in microfluidic oil droplets, after which they were lysed and their transcriptomes amplified, sequenced, and reads mapped, followed by data analysis. b) Barplot showing number of sequenced cells prepared by FACS sorting from each organ (n=20). c) Barplot showing number of sequenced cells prepared by microfluidic droplets from each organ (n=12).

We developed a procedure to collect 20 organs and tissues from the same mouse in which aorta, bladder, bone marrow, brain (cerebellum, cortex, hippocampus, striatum), diaphragm, fat (brown, gonadal, mesenteric, subcutaneous), heart, kidney, large intestine, limb muscle, liver, lung, mammary gland, pancreas, skin, spleen, thymus, tongue, and trachea were immediately dissected and processed into single cell suspensions, which in turn were either single cell sorted into plates with FACS or loaded into microfluidic droplets (see Extended Data and Methods). Single cell transcriptomes were sequenced to an average depth of 814,488 reads per cell for the plate data and 7,709 unique molecular identifiers (UMI) per cell for the microfluidic droplet data. After quality control filtering, 44,949 FACS sorted cells and 55,656 microfluidic droplet processed cells were retained for further analysis. A comparison of the two methods shows differences for each organ in the number of cells analyzed (Fig. 1b,c), reads per cell (Supp. Fig. 1a,c) and genes per cell (Supp. Fig. 1b,d).

We performed unbiased graph-based clustering of the pooled set of transcriptomes across all organs, and visualized them using tSNE (Fig. 2 and Supp. Fig. 2). The majority of clusters contain cells from only one organ (n=29/54), but a number of clusters (n=25/54) (Supp. Fig. 2) contained cells from multiple organs. To further dissect these clusters we analyzed each organ independently, first by performing principal component analysis (PCA) on the most variable genes in the organ, followed by nearest-neighbor graph-based clustering. We then used cluster-specific gene expression of known markers as well as genes differentially expressed between clusters to assign cell type annotations (Fig. 3, Supp.Fig.3, TableS1). A detailed description of the cell types and defining genes for each organ and tissue is available in the Supplementary Information. We used a standardized analysis approach for all organs and tissues and an example using liver can be found in the Organ Annotation Vignette. For each cell, we provide annotations in the controlled vocabulary of a cell ontology^19^ to facilitate comparisons with other experiments. Many of these cell clusters have not previously been obtained in pure populations and our data provide a wealth of new information on their characteristic gene expression profiles. Initial annotation of the cellular diversity of each organ and tissue can be found in the extended data, and a detailed discussion of each cell type on an organ by organ basis can be found in the supplement. Some unexpected discoveries include a potential new role for genes *Neurog3, Hex3,* and *Prss53* in the adult pancreas, a cell population expressing *Chodl* in limb muscle, transcriptional heterogeneity of brain endothelial cells, the expression of MHCII genes by adult mouse T cells, and sets of transcription factors that can specifically distinguish between similar cell types across multiple organs and tissues.

**Figure 2.**
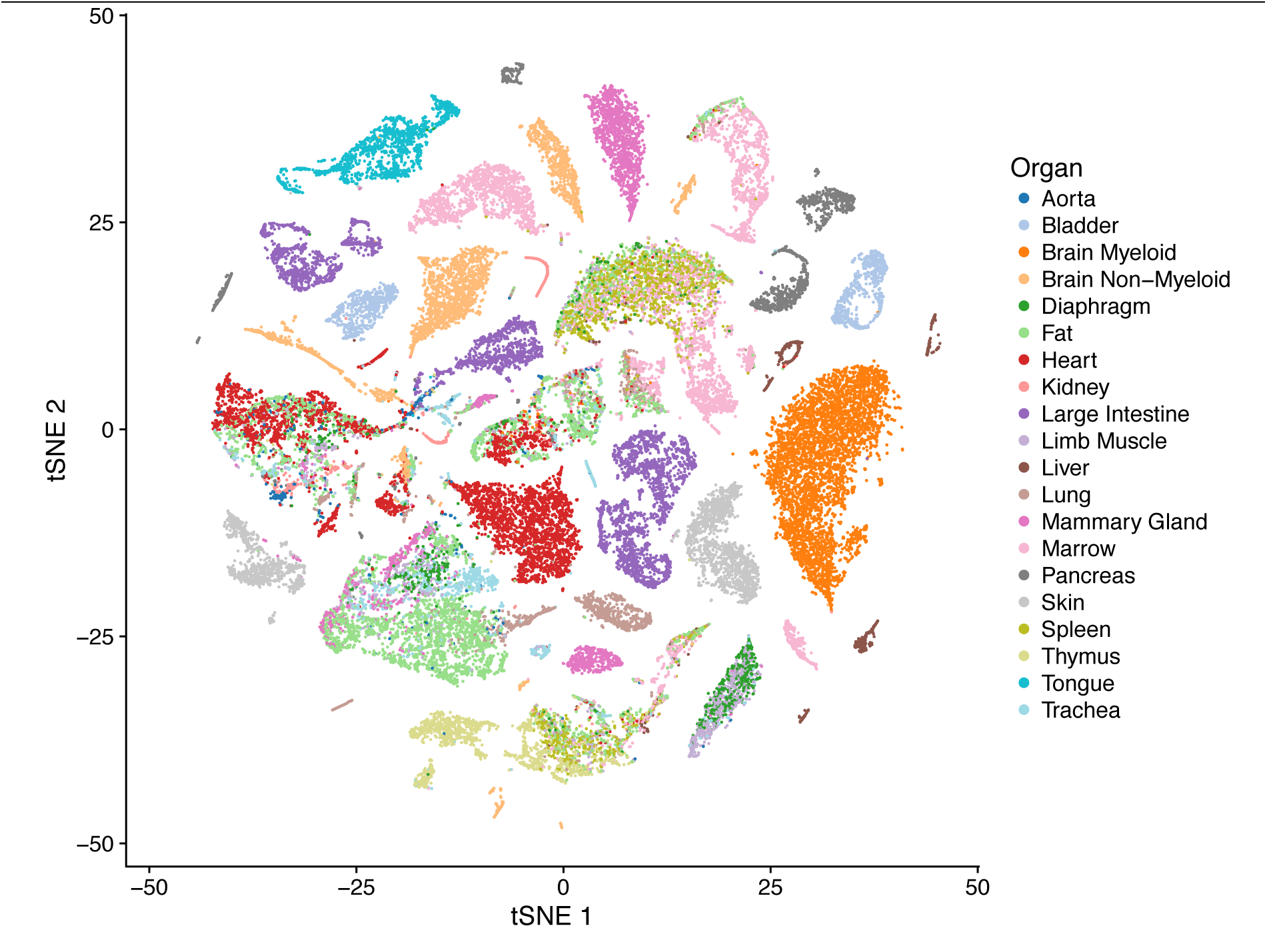
tSNE visualization of all FACS sorted cells. tSNE plot of all cells sorted by FACS, color coded by organ.

**Figure 3.**
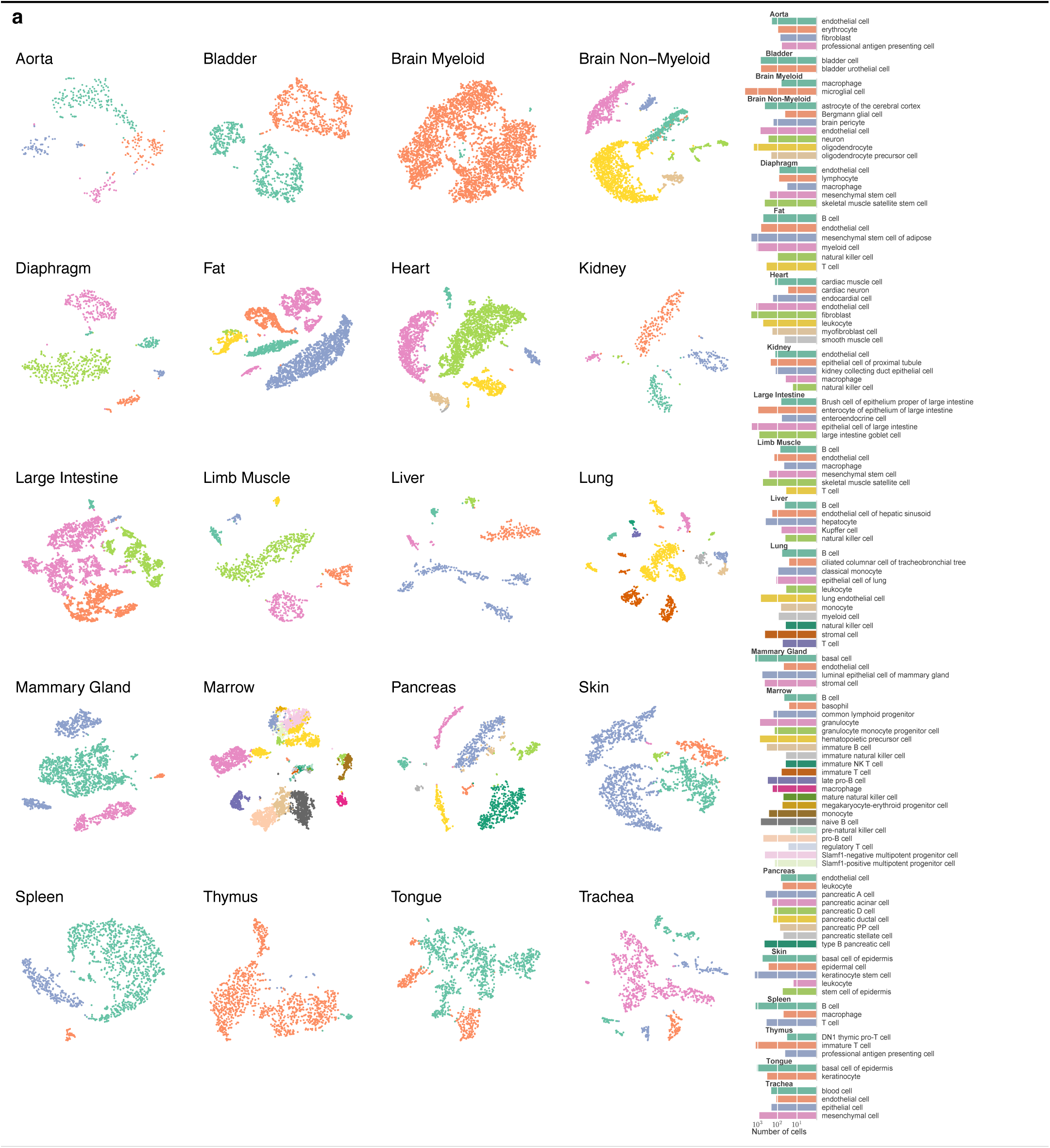
tSNE visualization of individual organs. a) tSNE plots for each organ of cells sorted by FACS. Color coding indicates distinct clusters. b) Barplots of annotated cell types based on differential gene expression across all organs. Coloring of clusters within each organ is consistent between panels a and b.

Any individual single-cell sequencing experiment offers a partial view of the diversity of cell types within an organism and the gene expression within each cell type. We illustrate the variability to be expected between methods and experiments by comparing our two measurement approaches to one another, and to data from Han *et al.*^20^ generated using a third method, microwell-seq. One striking feature is the variability in the number of genes detected per cell between organs and tissues and between methods. For example, the median number of genes detected per cell in bladder is about 4900 in the FACS data, 2900 in the droplet data, and 900 in the microwell-seq data, while the number detected in kidney is about 1400 in the FACS data, 1900 in the droplet data, and 500 in the microwell-seq data. The bladder, liver, lung, mammary gland, trachea, tongue, and spleen all show nearly twice as many genes detected per cell in the FACS data as compared to the microfluidic data, whereas heart and marrow show comparable numbers detected in both methods (Supp. Fig. 4a). This difference does not appear to be due to sequencing depth, as the microfluidic droplet libraries are nearly saturated (Supp. Fig. 4b) and deeper sequencing of the FACS libraries could only increase the number of genes detected. In every organ, there are fewer genes detected per cell in microwell-seq data than either droplet or FACS data. In these comparisons, a gene is considered detected if a single read maps to it, as that is the only standard for expression at which reads and UMIs can be treated equally. We also looked at how the number of detected genes across each organ changes with different thresholds on the number of reads or UMIs (Supp. Fig. 5). We found that the number of detected genes decreases monotonically with increasing thresholds at similar rates across different organs and tissues within each method. We observed that in the droplet data more than half of the detected genes are represented by only a single UMI; this is to be expected given that only a few thousand UMIs are captured per cell. The FACS data are sampled much more deeply and one needs to set a relatively high threshold of 40 reads to see a comparable reduction in gene detection sensitivity.

**Figure 4.**
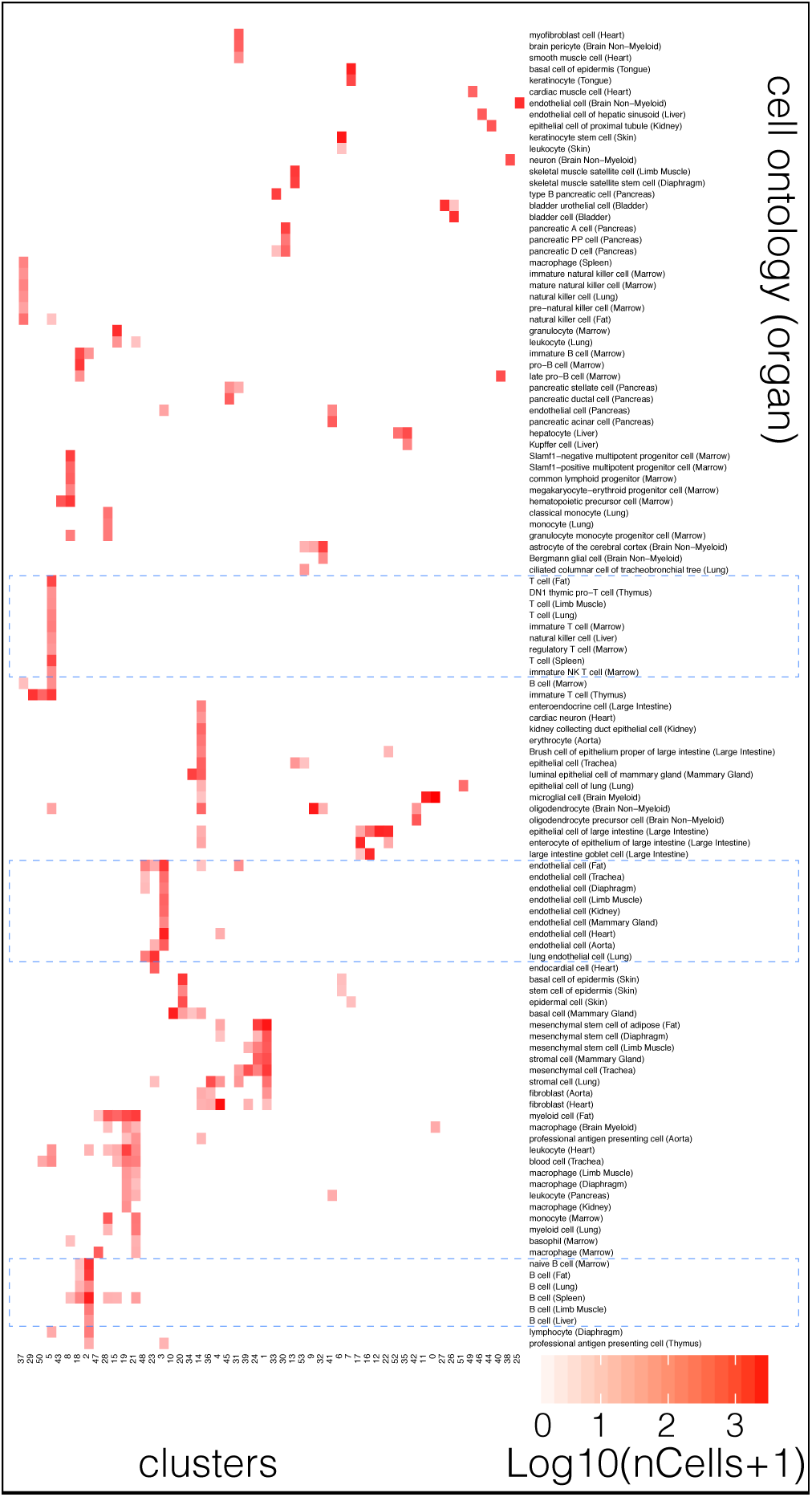
Comparison of cell type determination. Comparison of cell type determination as done by unbiased whole transcriptome comparison versus manual annotation by organ-specific experts. The x-axis represents clusters from Figure 2 and Figure S2 with multiple organs contributing, while the y-axis represents manual expert annotation of cell types in an organ-specific fashion. The unbiased method discovers relationships between similar cell types found in different organs (highlighted regions); in particular it groups T cells from different organs into a single cluster, B cells from different organs into a different single cluster, and endothelial cells from different organs into a single cluster.

**Figure 5.**
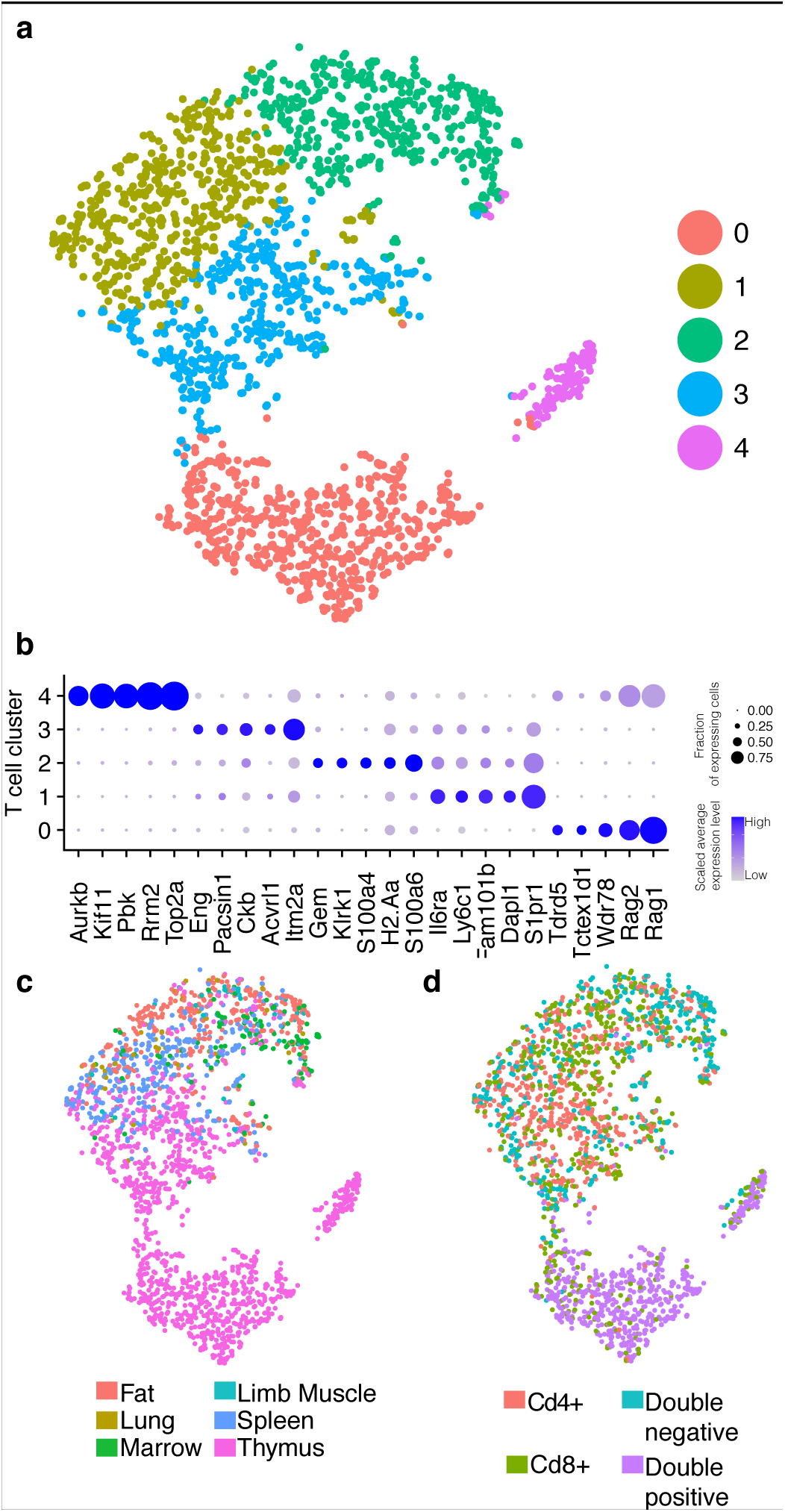
Analysis of all sorted T-cells. a) tSNE plot of all T cells colored by cluster membership. Five clusters were identified. b) Dotplot showing level of expression (color scale) and number of expressing cells (point diameter) within each cluster of T cells. c) tSNE plot of all T cells colored by organ of origin (Fat, Lung, Marrow, Limb Muscle, Spleen or Thymus). d) tSNE plot of all T cells colored by classification of T cells to 4 categories based on expression of Cd4 and Cd8 (Cd4^+^/ Cd8^+^/ Cd4^+^Cd8^+^ / Cd4^-^Cd8^-^).

Next, we investigated whether the three methods demonstrate concordance on the genes which define each of the cell clusters. To do so, we computed lists of genes (see Methods “Differential expression overlap analysis”) that differentiate between each cell cluster and the rest of the cell clusters in each organ across all three methods, focusing on common organs and cell clusters for the three methods. As expected, data from FACS and microfluidic droplet are in better agreement due to the fact that cells originated from the exact same organ or tissue and were prepared in parallel. For each cell cluster there appears to be a core of a few hundred defining genes on which all three methods agree (Supp. Fig. 6 and Table S2). This comparison suggests that independent datasets generated from the various tissue atlases that are beginning to arise can be combined and collectively analyzed to generate more robust characterizations of gene expression.

**Figure 6.**
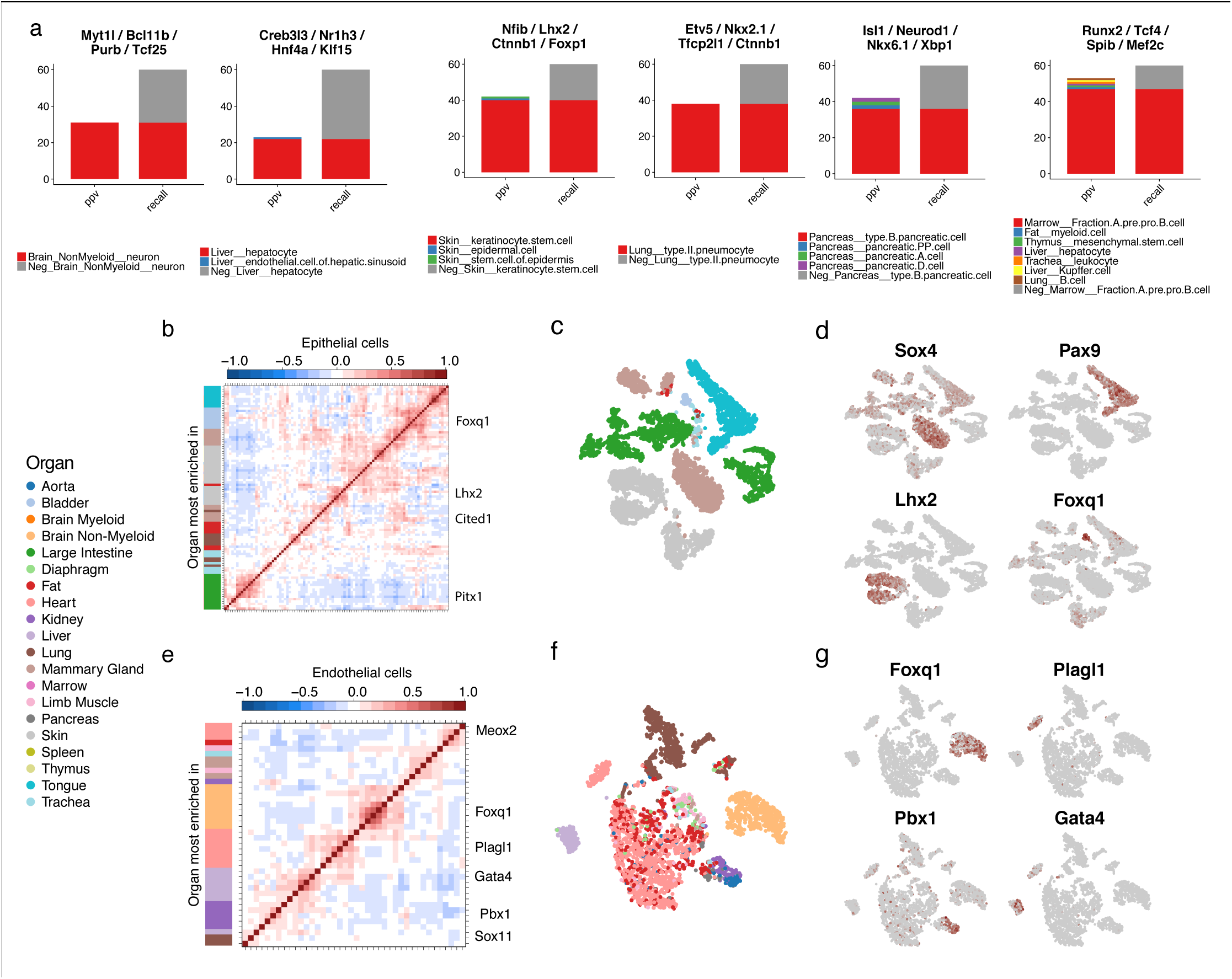
Transcription factor (TF) expression analysis. a) Visualization of the precision (ppv) and recall of combinations of 4 TFs. Red bars indicate the number of cells expressing all 4 TFs in the target cell type (true positive) in both the ppv and recall columns. Other colored bars in the ppv column represent the number of cells in the non-target cell types expressing all 4 TFs (false positives). The height of the grey bar in the recall column is the number of cells in the target cell type not expressing all 4 TFs (false negatives). The legend indicates the target cell type next to the red square and all non-target cell types with coexpression. Data shown is the entire dataset subsampled to at most 60 cells per cell type. b) Correlogram of top organ-specific TFs for epithelial cells. Row colors correspond to organ of the most-enriched cell type. c) tSNE visualization of epithelial cells, colored by organ. d) tSNE visualization of endothelial cell expression of select TFs. (grey/low to red/high). e) Correlogram of top organ-specific TFs for epithelial cells. Row colors correspond to organ of the most-enriched cell type. f) tSNE visualization of epithelial cells, colored by organ. g) tSNE visualization of epithelial cell expression of select TFs.

To understand the relationships between cell types, we mapped the annotations of organ-specific cell types onto the unbiased clustering of all cells. It is evident that the clusters in Figure 2 (also Supp. Fig. 2) containing cells from multiple organs generally represent shared cell types common to those organs (Fig. 4). For example, B cells from fat, limb muscle, diaphragm, lung, spleen and marrow cluster together, as do T cells from spleen, marrow, lung, limb muscle, fat and thymus. Interestingly, while endothelial cells from fat, heart, and lung cluster together, they are segregated from endothelial cells from the mammary gland, kidney, trachea, limb muscle, aorta, diaphragm, and pancreas. Such differences could be caused by true differential gene expression signatures across different organs, but could also potentially be influenced by organ-specific batch effects. The fact that many cells cluster together across organs and biological replicates is evidence that batch effects are not the main source of variance in the dataset. Our findings show that manual annotation of cell types is consistent with unbiased transcriptomic clustering, and that most cell types are unique enough to enable their unbiased identification across organs and tissues. We expect that further refinements of comparison algorithms will facilitate the discovery of finer, organ-specific distinctions between these shared cell types.

To investigate common cell types across all organs, we pooled all cells annotated as T cells and analyzed them collectively (Fig. 5). Our analysis revealed 5 clusters. Cluster 0 comprises cells from the thymus that are undergoing VDJ recombination characterized by the expression of RAG (*Rag1, Rag2*) and TdT (*Dntt*), and includes uncommitted double positive T-cells (*Cd4^+^, Cd8a*^*+*^). Cluster 4 contains proliferating T cells, predominantly from the thymus. We hypothesize that these are pre-T cells expanding after the completion of VDJ recombination. Clusters 1-3 contain predominantly single positive T cells (*Cd4*^*+*^ or *Cd8a*^+^). Cluster 3 contains *Cd5*^high^ thymic T cells possibly undergoing positive selection while Cluster 2 contains mostly non-thymic T cells expressing the high affinity IL2 receptor (*Il2ra, Il2rb*), suggesting they are activated. Interestingly, they also express MHC type II genes (*H2-Aa, H2-Ab1*). While this is known to occur in human T cells, MHCII was previously thought restricted to professional antigen presenting cells in mice^11^. Finally, Cluster 1 also represents mature T cells, but primarily from the spleen.

A key challenge for many single cell studies is understanding the potential changes to the transcriptome caused by handling, dissociation and other experimental manipulation. A previous study in limb muscle showed that quiescent satellite cells tend to become activated by dissociation and consequently express immediate early genes among other genes^21^. We found that expression of these dissociation-related markers was also clearly observed in our limb muscle data, as well as in mammary gland and bladder (Supp. Fig. 7), but that many organs and tissues showed little evidence of similar cellular activation. Therefore the dissociation-related activation markers found in limb muscle are not universal across all organs and tissues. This is not to say that other organs lack dissociation-related gene expression changes, but that some of the genes involved are specific to a given organ. Importantly, the presence of such gene expression changes does not prevent the identification of cell type or the comparison of cell types across organs and tissues.

One major goal of defining cell identities is to understand the transcription factor (TF) regulatory networks that underlie them. We first investigated the combinatorial specificity of TF expression across all cell types (defined as unique combinations of cell ontology annotation and tissue) (**Fig. 6**). We searched for the combination of four (n=4) enriched TFs that best specified each target cell type over all others. For each combination of TFs, we counted every cell expressing all four TFs as a positive, and anything else as a negative. We then calculated cell type-specificity by the precision (ratio of number of positive target cells to total number of positive cells) and recall (ratio of number of positive target cells to total number of target cells) of each combination of TFs for the target cell type over the rest of the cells (**Table S3**). We found 41 cell types with TF combinations with precision > 0.3 and recall > 0.3. We noted that the combinatorial nature of TF expression was critical to specificity; for example, *Ctnnb1,* combined with one of two TF sets, specified either skin keratinocyte stem cells or lung type II pneumocytes (**Fig. 6a**). We found many TF combinations for cell types with challenging *in vitro* differentiation protocols (e.g., hepatocytes; *Creb3l3, Nr1h3, Hnf4a,* and *Klf15*) and cell types with no established direct differentiation protocol (e.g., microglia; *Mafb, Sall1, Irf5,* and *Maf*) (**Fig. 6a**).

We then analyzed organ-specific TFs by isolating a set of closely-related, cross-organ cell groups (epithelial cells and endothelial cells). We performed TF correlation analysis, similar to ^15^ within the cell groups (**Fig. 6b-g**). We found many TFs within epithelial cells that clustered strongly by organ and were enriched in organ-specific epithelial clusters (**Fig. 6b**). For example, *Sox4* (mammary basal cells), *Foxq1* (bladder basal cells of the urothelium), *Pax9* (tongue basal cells of the epidermis), and *Lhx2* (skin keratinocyte stem cells) were highly organ-specific (**Fig. 6c,d**). Within endothelial cells, liver, brain, mammary gland/limb muscle, and lung-specific clusters of TFs were evident (**Fig. 6e-g**). *Gata4,* known to specify liver endothelium, appeared in a cluster of liver-enriched TFs (**Fig. 6g**). Another cluster of TFs, including *Pbx1,* were enriched in kidney endothelial cells (**Fig. 6g**). The roles of *Pbx1* in kidney endothelial development are not explored, and could aid in tissue engineering for kidney regeneration. A highly distinct cluster of cells specified the heart endocardium, including *Plagl1,* a TF whose role in endocardial specification is unknown (**Fig. 6g**). These results illustrate how single cell data taken across many organs and organs can identify the transcriptional regulatory programs which are specific to cell types of interest.

In conclusion, we have created a compendium of single-cell transcriptional measurements across 20 organs and tissues of the mouse. This *Tabula Muris,* or “Mouse Atlas”, has many uses, including the discovery of new putative cell types, the discovery of novel gene expression in known cell types, and the ability to compare cell types across organs and tissues. It will also serve as a reference of healthy young adult organs and tissues which can be used as a baseline for current and future mouse models of disease. While it is not an exhaustive characterization of all organs of the mouse, it does provide a rich data set of the most highly studied organs and tissues in biology. The *Tabula Muris* provides a framework and description of many of the most populous and important cell populations within the mouse, and represents a foundation for future studies across a multitude of diverse physiological disciplines.

**Supplementary Information** is available in the online version of the paper.

## Acknowledgements

We thank Sony Biotechnology for making an SH800S instrument available for this project. Some cell sorting/flow cytometry analysis for this project was done on a Sony SH800S instrument in the Stanford Shared FACS Facility. Some fluorescence activated cell sorting (FACS) was done with instruments in the VA Flow Cytometry Core, which is supported by the US Department of Veterans Affairs (VA), Palo Alto Veterans Institute for Research (PAVIR), and the National Institutes of Health (NIH).

## Methods

### Mice and Tissue Collection

Four 10-15 week old male and four virgin female C57BL/6 mice were shipped from the National Institute on Aging colony at Charles River to the Veterinary Medical Unit (VMU) at the VA Palo Alto (VA). At both locations, mice were housed on a 12-h light/dark cycle, and provided food and water *ad libitum.* The diet at Charles River was NIH-31, and Teklad 2918 at the VA VMU. Littermates were not recorded or tracked, and mice were housed at the VA VMU for no longer than 2 weeks before euthanasia. Prior to tissue collection, mice were placed in sterile collection chambers for 15 minutes to collect fresh fecal pellets. Following anesthetization with 2.5% v/v Avertin, mice were weighed, shaved, and blood drawn via cardiac puncture before transcardial perfusion with 20 ml PBS. Mesenteric adipose tissue (MAT) was then immediately collected to avoid exposure to the liver and pancreas perfusate, which negatively impacts cell sorting. Isolating viable single cells from both pancreas and liver of the same mouse was not possible, therefore, 2 males and 2 females were used for each. Whole organs were then dissected in the following order: large intestine, spleen, thymus, trachea, tongue, brain, heart, lung, kidney, gonadal adipose tissue (GAT), bladder, diaphragm, limb muscle *(tibialis anterior),* skin (dorsal), subcutaneous adipose tissue (SCAT, inguinal pad), mammary glands (fat pads 2, 3, and 4), brown adipose tissue (BAT, interscapular pad), aorta, and bone marrow (spine and limb bones). Following single cell dissociation as described below, cell suspensions were either used for FACS sorting of individual cells into 384-well plates, or for microfluidic droplet library preparation. All animal care and procedures were carried out in accordance with institutional guidelines approved by the VA Palo Alto Committee on Animal Research.

### Tissue dissociation and sample preparation

Specific protocols for each tissue are described in the supplement.

## Single Cell Methods

### Lysis plate preparation

Lysis plates were created by dispensing 0.4 μl lysis buffer (0.5 U Recombinant RNase Inhibitor (Takara Bio, 2313B), 0.0625% Triton™ X-100 (Sigma, 93443-100ML), 3.125 mM dNTP mix (Thermo Fisher, R0193), 3.125 μM Oligo-dT_30_VN (IDT, 5’AAGCAGTGGTATCAACGCAGAGTACT**30**VN-3’) and 1:600,000 ERCC RNA spike-in mix (Thermo Fisher, 4456740)) into 384-well hard-shell PCR plates (Biorad HSP3901) using a Tempest liquid handler (Formulatrix). 96-well lysis plates were also prepared with 4 μl lysis buffer. All plates were sealed with AlumaSeal CS Films (Sigma-Aldrich Z722634) and spun down (3,220 x g, 1 minute) and snap frozen on dry ice. Plates were stored at −80°C until sorting.

### FACS sorting

After dissociation, single cells from each organ and tissue were isolated into 384- or 96-well plates via Fluorescence Activated Cell Sorting (FACS). Most organs were sorted into 384-well plates using SH800S (Sony) sorters. Heart and liver were sorted into 96-well plates and cardiomyocytes were hand-picked into 96-well plates. Limb muscle and diaphragm were sorted into 384-well plates on an Aria III (Becton Dickinson) sorter. The last two columns of each 384 well plate were intentionally left as blanks. For most organs, single cells were selected with forward scatter, and dead cells and common cell types were excluded with a single color channel. Combinations of fluorescent antibodies were used for most organs to enrich for rare cell populations (see supplemental text), but some were stained only for viable cells. Color compensation was used whenever necessary. On the SH800, the highest purity setting (“Single cell”) was used for all but the rarest cell types, for which the “Ultrapure” setting was used. Sorters were calibrated using FACS buffer every day before collecting any cells, and also after every 8 sorted plates. For a typical sort, 1-3 ml of pre-stained cell suspension was filtered, vortexed gently, and loaded onto the FACS machine. A small number of cells were flowed at low pressure to check cell and debris concentrations. The pressure was then adjusted, flow paused, the first destination plate unsealed, loaded and sorting started. If a cell suspension was too concentrated, it was diluted using FACS buffer or 1X PBS. For some cell types like hepatocytes, 96-well plates were used because it was not possible to sort individual cells accurately into 384-well plates. Immediately after sorting, plates were sealed with a pre-labeled aluminum seal, centrifuged, and flash frozen on dry ice. On average, each 384-well plate took 8 minutes to sort.

### cDNA synthesis and library preparation

cDNA synthesis was performed using the Smart-seq2 protocol^2,3^. Briefly, 384-well plates containing single-cell lysates were thawed on ice followed by first strand synthesis. 0.6 μl of reaction mix (16.7 U/μl SMARTScribe Reverse Transcriptase (Takara Bio, 639538), 1.67 U/μl Recombinant RNase Inhibitor (Takara Bio, 2313B), 1.67X First-Strand Buffer (Takara Bio, 639538), 1.67 μM TSO (Exiqon, 5’-AAGCAGTGGTATCAACGCAGAGTGAATrGrGrG-3’), 8.33 mM DTT (Bioworld, 40420001-1), 1.67 M Betaine (Sigma, B0300-5VL), and 10 mM MgCl_2_ (Sigma, M1028-10X1ML)) was added to each well using a Tempest liquid handler. Reverse transcription was carried out by incubating wells on a ProFlex 2 x 384 thermal-cycler (Thermo Fisher) at 42°C for 90 minutes, and stopped by heating at 70°C for 5 minutes.

Subsequently, 1.5 μl of PCR mix (1.67X KAPA HiFi HotStart ReadyMix (Kapa Biosystems, KK2602), 0.17 μM IS PCR primer (IDT, 5’-AAGCAGTGGTATCAACGCAGAGT-3’), and 0.038 U/μl Lambda Exonuclease (NEB, M0262L)) was added to each well with a Mantis liquid handler (Formulatrix), and second strand synthesis was performed on a ProFlex 2×384 thermal-cycler by using the following program: 1) 37°C for 30 minutes, 2) 95°C for 3 minutes, 3) 23 cycles of 98°C for 20 seconds, 67°C for 15 seconds, and 72°C for 4 minutes, and 4) 72°C for 5 minutes.

The amplified product was diluted with a ratio of 1 part cDNA to 10 parts 10mM Tris-HCl (Thermo Fisher, 15568025), and concentrations were measured with a dye-fluorescence assay (Quant-iT dsDNA High Sensitivity kit; Thermo Fisher, Q33120) on a SpectraMax i3x microplate reader (Molecular Devices). Sample plates were selected for downstream processing if the mean concentration of blanks (ERCC-containing, non-cell wells) was greater than 0 ng/μl, and, after linear regression of the values obtained from the Quant-iT dsDNA standard curve, the R^2^ value was greater than 0.98. Sample wells were then selected if their cDNA concentrations were at least one standard deviation greater than the mean concentration of the blanks. These wells were reformatted to a new 384-well plate at a concentration of 0.3 ng/μl and final volume of 0.4 μl using an Echo 550 acoustic liquid dispenser (Labcyte).

Illumina sequencing libraries were prepared as described in Darmanis et al. 2015.^4^ Briefly, tagmentation was carried out on double-stranded cDNA using the Nextera XT Library Sample Preparation kit (Illumina, FC-131-1096). Each well was mixed with 0.8 μl Nextera tagmentation DNA buffer (Illumina) and 0.4 μl Tn5 enzyme (Illumina), then incubated at 55°C for 10 minutes. The reaction was stopped by adding 0.4 μl “Neutralize Tagment Buffer” (Illumina) and centrifuging at room temperature at 3,220 x g for 5 minutes. Indexing PCR reactions were performed by adding 0.4 μl of 5 μM i5 indexing primer, 0.4 μl of 5 μM i7 indexing primer, and 1.2 μl of Nextera NPM mix (Illumina). PCR amplification was carried out on a ProFlex 2×384 thermal cycler using the following program: 1) 72°C for 3 minutes, 2) 95°C for 30 seconds, 3) 12 cycles of 95°C for 10 seconds, 55°C for 30 seconds, and 72°C for 1 minute, and 4) 72°C for 5 minutes.

### Library pooling, quality control, and sequencing

Following library preparation, wells of each library plate were pooled using a Mosquito liquid handler (TTP Labtech). Pooling was followed by two purifications using 0.7x AMPure beads (Fisher, A63881). Library quality was assessed using capillary electrophoresis on a Fragment Analyzer (AATI), and libraries were quantified by qPCR (Kapa Biosystems, KK4923) on a CFX96 Touch Real-Time PCR Detection System (Biorad). Plate pools were normalized to 2 nM and equal volumes from 10 or 20 plates were mixed together to make the sequencing sample pool. A PhiX control library was spiked in at 0.2% before sequencing.

### Sequencing libraries from 384-well and 96-well plates

Libraries were sequenced on the NovaSeq 6000 Sequencing System (Illumina) using 2 x 100bp paired-end reads and 2 x 8bp or 2 x 12bp index reads with either a 200- or 300-cycle kit (Illumina, 20012861 or 20012860).

### Microfluidic droplet single cell analysis

Single cells were captured in droplet emulsions using the GemCode Single-Cell Instrument (10x Genomics, Pleasanton, CA, USA), and SC RNA-seq libraries were constructed as per the 10X Genomics protocol using GemCode Single-Cell 3’ Gel Bead and Library V2 Kit. Briefly, single cell suspensions were examined using an inverted microscope, and if sample quality was deemed satisfactory, the sample was diluted in PBS with 2% FBS to a concentration of 1000 cells/μl. If cell suspensions contained cell aggregates or debris, two additional washes in PBS with 2% FBS at 300 x g for 5 minutes at 4°C were performed. Cell concentration was measured either with a Moxi GO II (Orflo Technologies) or a hemocytometer. Cells were loaded in each channel with a target output of 5,000 cells per sample. All reactions were performed in the Biorad C1000 Touch Thermal cycler with 96-Deep Well Reaction Module. 12 cycles were used for cDNA amplification and sample index PCR. Amplified cDNA and final libraries were evaluated on a Fragment Analyzer using a High Sensitivity NGS Analysis Kit (Advanced Analytical). The average fragment length of 10x cDNA libraries was quantitated on a Fragment Analyzer (AATI), and by qPCR with the Kapa Library Quantification kit for Illumina. Each library was diluted to 2 nM, and equal volumes of 16 libraries were pooled for each NovaSeq sequencing run. Pools were sequenced with 100 cycle run kits with 26 bases for Read 1, 8 bases for Index 1, and 90 bases for Read 2 (Illumina 20012862). A PhiX control library was spiked in at 0.2 to 1%. Libraries were sequenced on the NovaSeq 6000 Sequencing System (Illumina)

### Data Processing

Sequences from the Novaseq were de-multiplexed using bcl2fastq version 2.19.0.316. Reads were aligned using to the mm10plus genome using STAR version 2.5.2b with parameters TK. Gene counts were produced using HTSEQ version 0.6.1p1 with default parameters, except “stranded” was set to “false”, and “mode” was set to “intersection-nonempty”.

Sequences from the microfluidic droplet platform were de-multiplexed and aligned using CellRanger, available from 10x Genomics with default parameters.

### Clustering

Standard procedures for filtering, variable gene selection, dimensionality reduction, and clustering were performed using the Seurat package. A detailed worked example, including the mathematical formulae for each operation, is in the Tissue Annotation Vignette. The parameters that were tuned on a per-tissue basis (resolution and number of PCs can be viewed in the tissue-specific Rmd files available on GitHub). For each tissue and each sequencing method (FACS and microfluidic droplet), the following steps were performed:

1. Cells were lexicographically sorted by cell ID to ensure reproducibility.
2. Cells with fewer than 500 detected genes were excluded. (A gene counts as detected if it has at least one read mapping to it). Cells with fewer than 50,000 reads (FACS) or 1000 UMI (microfluidic droplet) were excluded.
3. Counts were log-normalized for each cell using the natural logarithm of 1 + counts per million (for FACS) or 1 + counts per ten thousand (for microfluidic droplet).
4. Variable genes were selected using a threshold (0.5) for the standardized log dispersion, where the standardization was done in separately according to binned values of log mean expression.
5. The variable genes were projected onto a low-dimensional subspace using principal component analysis. The number of principal components was selected based on inspection of the plot of variance explained.
6. A shared-nearest-neighbors graph was constructed based on the Euclidean distance in the low-dimensional subspace spanned by the top principal components. Cells were clustered using a variant of the Louvain method that includes a resolution parameter in the modularity function^23^.
7. Cells were visualized using a 2-dimensional t-distributed Stochastic Neighbor Embedding of the PC-projected data.
8. Cell types were assigned to each cluster using the abundance of known marker genes. Plots showing the expression of the markers for each tissue appear in the extended data.
9. When clusters appeared to be mixtures of cell types, they were refined either by increasing the resolution parameter for clustering or subsetting the data and rerunning steps 3-7.

A similar analysis was done globally for all FACS processed cells and for all microfluidic droplet processed cells to produce an unbiased clustering.

### Differential expression overlap analysis

For FACS and microfluidic droplet data differential expression analysis for each organ was performed using a Wilcox rank test as implemented in the “FindAllMarkers” function of the Seurat package. Differential expression was performed between cell ontology groups and resulted in a list of differentially expressed genes (log_e_FoldChange > 0.25) between each cell ontology group and all other ontology groups of the same organ. For the microwellSeq we used the corresponding published lists for each cell type and for every organ. We then assessed the overlap (Supp. Fig. 6) of those lists between the three methods. As the nomenclature is not identical, the analysis was performed between cell types that could be matched with a certain degree of confidence between the three methods (TableS2).

### Calculation of dissociation scores

For each organ, gene expression matrices were subset to 140 genes, and principal component analysis was performed on this gene subset. The first principal component was used as the “dissociation score” as it corresponds to the variance within these genes.

### Defining cell type-enriched transcription factors

Transcription factors were defined as the 1140 genes annotated by the Gene Ontology term “DNA binding transcription factor activity”, downloading from the Mouse Genome Informatics database (http://www.informatics.jax.org/mgihome/GO/project.shtml, accessed on 2017-11-10). Cell types were defined as unique combinations of cell ontology and organ annotation (e.g. Lung__Endothelial_cell). All analysis was performed on the full 3 month dataset, subsampled by randomly selecting 60 cells from each cell type. Enriched TFs were defined by the Seurat FindMarkers function with the “Wilcoxon” significance test for the target cell type against the all of rest of the cell types combined. These were filtered by p_val < 10-3, avg_diff > 0.2, pct.1 – pct.2 > 0.1 (percent detected difference > 0.1), and pct.1 > 0.3 (detected in > 30% of target cells).

### Discovering cell type-specific TF combinations

For each cell type that contained at least 6 cells, and had at least 4 enriched TFs, the top 30 TFs or all that passed filter, whichever was smaller, were selected by highest avg_diff. The specificity of each four-TF combination (up to 27405 combinations for 30 TFs) was assessed by a score defined from two standard metrics, precision and recall:

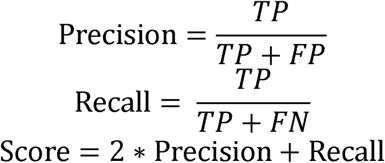

Where TP (true positive) is the number of cells in the target cell type expressing all 4 TFs, FP (false positive) is the number of cells not in the target cell type expressing all 4 TFs, and TN (true negative) is the number of cells in the target cell type not expressing all 4 TFs. The top TFs by this score for several cell types was plotted in Figure 6a.

### Defining TF networks by correlation analysis

Organ-specific TF regulatory networks were measured by the correlations of TFs. TFs were selected by enrichment in a cell type over all other cell type with the test described in “Defining cell type-enriched transcription factors”, filtered by p_val < 10, avg_diff > 0.3, and pct.1-pct.2 > 0.1. The top 8 markers per cell type (or however many passed the filters) were selected by avg_diff. The Pearson correlations between genes were calculated, and genes ordered by hierarchical clustering with optimal ordering (hclust and cba::optimal). For analysis of TFs within single broad cross-organ cell types, endothelial cells were defined as cell ontology annotations containing “endothelial” or “capillary” (Fig. 6e-g). Epithelial cells were defined as cell ontology annotations containing “epithelial”, “basal”, “keratinocyte”, or “epidermis” (Fig. 6b-d). Exemplary organ-specific TFs were visualized on t-SNE plots. t-SNE was computed for a single cell annotation across all organs, by the top variable genes (Seurat FindVariableGenes, RunPCA with 10 PCs, and RunTSNE with perplexity = 30).

## Supplementary Figure Captions

**Supplementary Figure 1** a) Histogram of number of reads per cell for each organ from FACS sorted cells. b) Histogram of number of genes detected per cell for each organ from FACS sorted cells. c) Histogram of number of unique molecular identifiers (UMI) sequenced per cell for each organ from cells prepared by microfluidic droplets. d) Histogram of number of genes detected per cell for each organ for cells prepared by microfluidic droplets.

**Supplementary Figure 2**. tSNE visualization of all FACS sorted cells annotated by cluster. Clusters are discussed in the text and further analyzed in Figure 4.

**Supplementary Figure 3** a) tSNE plot of all cells captured by microfluidic droplets color coded by organ. b) Dimensionally reduced tSNE plots for each organ of cells sorted by microfluidic droplets. Color coding indicates distinct clusters. c) Barplots of manually annotated cell types based on differential gene expression across all organs. Coloring of clusters within each organ is consistent between panels b and c.

**Supplementary Figure 4** a) Number of genes detected by FACS (red), microfluidic droplets (green) and microwell-Seq (blue) (Han *et al*.). b) library saturation fraction for all 10x libraries included in the study. Dotted horizontal line demarcates the median (=0.86).

**Supplementary Figure 5** Fraction of all detectable genes, for each cell across all organs, (UMI/read threshold is >0) detected at increasing UMI/read thresholds for FACS (left), microfluidic droplet (middle) and microwell-Seq (right).

**Supplementary Figure 6** Venn diagrams showing the overlap between differentially expressed genes for each common cell type and organs across three methods (FACS, droplet, microwell-Seq). Plotted data are provided in tabular form in Table S2.

**Supplementary Figure 7** Analysis of dissociation induced gene expression scores across organs.

## Supplementary Tables

**Supplementary Table 1** Number of cells belonging to each annotated cell type across all organs for FACS and microfluidic droplets.

**Supplementary Table 2** Cell type comparisons and lists of differentially expressed genes across three methods (FACS, droplet, microwell-Seq) and all common organs and tissues.

**Supplementary Table 3** Combinatorial specificity of transcription factors (TFs) to single cell types. Three combinations of 4 TFs with the highest combinatorial specificity score are presented. The precision (ppv) and recall of each 4-TF combination and cell type is calculated as described in the Methods and main text.

